# Simultaneous multiplexed amplicon sequencing and transcriptome profiling in single cells

**DOI:** 10.1101/328328

**Authors:** Mridusmita Saikia, Philip Burnham, Sara H. Keshavjee, Michael F. Z. Wang, Michael Heyang, Pablo Moral-Lopez, Meleana M. Hinchman, Charles G. Danko, John S. L. Parker, Iwijn De Vlaminck

## Abstract

We describe Droplet Assisted RNA Targeting by single cell sequencing (DART-seq), a versatile technology that enables multiplexed amplicon sequencing and transcriptome profiling in single cells. We applied DART-seq to simultaneously characterize the non-A-tailed transcripts of a segmented dsRNA virus and the transcriptome of the infected cell. In addition, we used DART-seq to simultaneously determine the natively paired, variable region heavy and light chain amplicons and the transcriptome of B lymphocytes.

---

Droplet microfluidics has made high-throughput single-cell RNA sequencing accessible to more laboratories than ever before, but is restricted to capturing information from the ends of A-tailed messenger RNA (mRNA) transcripts^1–3^. Here we report DART-seq, a method that enables high-throughput targeted RNA amplicon sequencing and transcriptome profiling in single cells. DART-seq achieves this via a simple and inexpensive alteration of the Drop-seq strategy^1^. Drop-seq relies on co-encapsulation of single cells with barcoded primer beads that capture and prime reverse transcription (RT) of cellular mRNA^1,2^. The primers on Drop-seq beads comprise a common PCR sequence, a bead-specific cell barcode, a unique molecular identifier (UMI), and a poly-dT sequence for mRNA RT priming. To enable simultaneous measurement of the transcriptome and multiplexed RNA amplicons in DART-seq, we devised a scheme to enzymatically attach custom primers to a subset of poly-dTs on Drop-seq beads. This is achieved by first annealing a double-stranded toehold probe with a 3’ ssDNA overhang that is complementary to the poly-dT sequence, and then ligating the toehold using T4 DNA ligase (Fig. S1). A variety of custom primers with different sequences can be attached to the same beads in a single reaction.

We characterized the efficiency, tunability and variability of the ligation reaction using fluorescence hybridization assays (Fig. 1a and Fig. S2). We found that the primer ligation reaction is highly efficient (25-40%), and the number of custom primers ligated to the beads is directly proportional to the number of primers included in the ligation reaction (Fig. 1a). This was true for four primer sequences tested over a wide range of primer concentrations. The efficiency of probe ligation decreased for ligation reactions with more than 10^10^ molecules per bead, indicating saturation of available oligo(dT)s. We compared the fluorescence hybridization signal across individual beads and found that the bead-to-bead variability in fluorescence signal is small (standard deviation 3.0%, Fig. 1b).

**Fig. 1:**
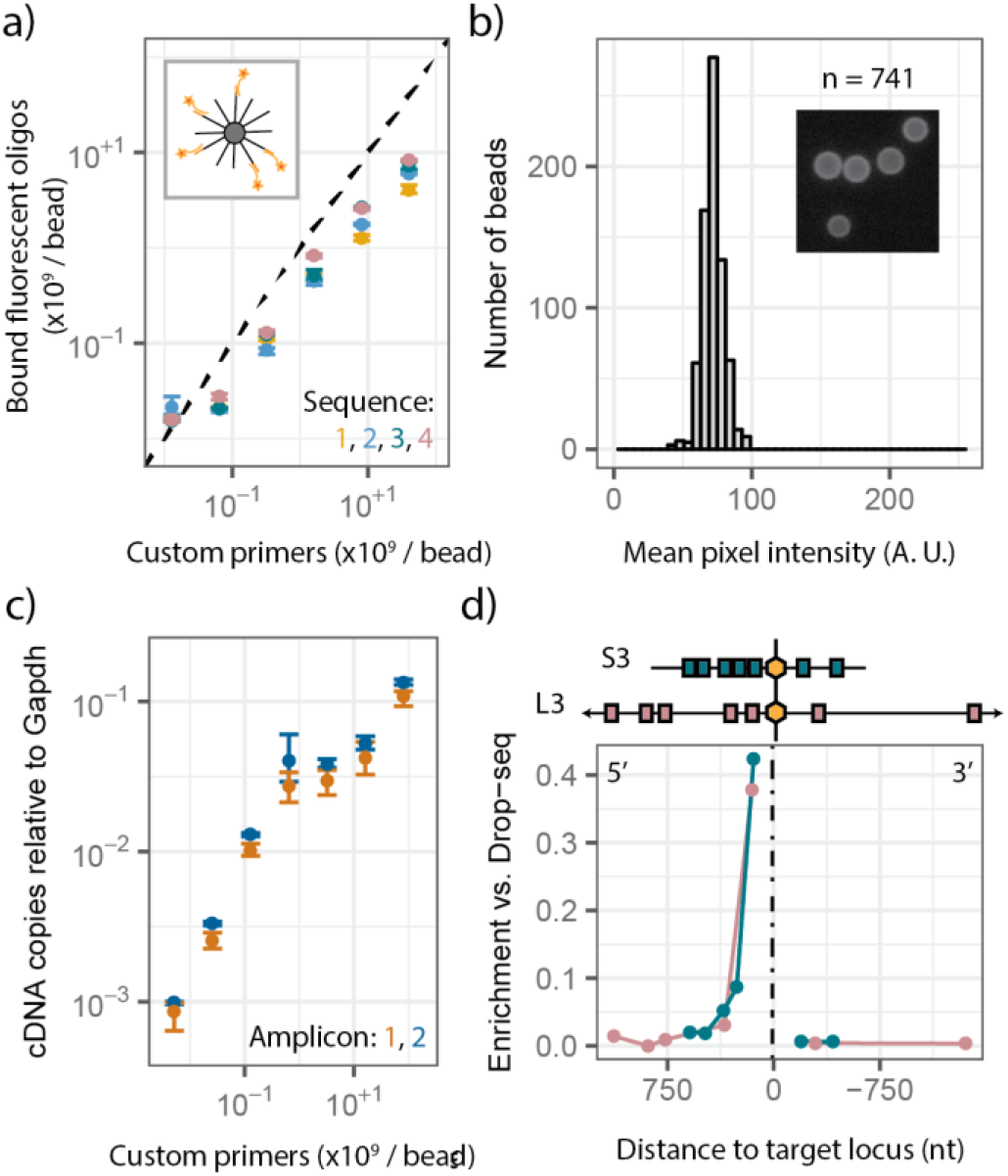
Characterization of DART-seq primer bead synthesis and RT priming. (a) Number of fluorescence probes bound per bead as function of the number of primers per bead included in the ligation reaction (four distinct primer sequences). Error bars indicate the minimum and maximum of three replicate measurements, points indicate the mean. The dotted line indicates expected values for 100% ligation efficiency. Inset: Schematic of fluorescence hybridization assay. (b) Bead-to-bead variability in fluorescence pixel intensity (n = 741 beads, maximum pixel intensity is 255). Inset: representative fluorescence microscopy image of beads. (c) cDNA copies of reovirus RNA relative to *Gapdh* as function of the number of custom primers included in the ligation reaction (bulk assay). Error bars indicate minimum and maximum of three replicates, points indicate the mean. (d) Enrichment of PCR amplicons relative to *Gapdh* in DART-seq versus Drop-seq libraries as function of distance to the target locus. Measurement for two reovirus genes (S3 in green and L3 in violet).

After primer bead synthesis, DART-seq follows the Drop-seq workflow without modification. Cells and primer beads are co-encapsulated in droplets using microfluidics. Cellular RNA is captured by the beads, and reverse transcribed. The DART-seq beads prime RT of both A-tailed mRNA and custom RNA amplicons. The resulting complementary DNA (cDNA) is PCR-amplified, tagmented, and again PCR amplified before sequencing. The cell-of-origin of mRNAs and RNA amplicons is identified by decoding the cell barcodes.

We assessed RT priming efficiency as a function of the number of custom primers ligated to DART-seq beads. We used quantitative PCR (qPCR) to measure the yield of cDNA copies of a non-A-tailed viral mRNA in reovirus-infected murine fibroblasts (L cells, Fig. 1c). Two distinct primers were ligated, targeting the same viral genome segment (S2). The yield of cDNA copies of viral mRNA, relative to cDNA copies of a host transcript (*Gapdh*), increased with the number of primers included in the ligation reaction, and saturated for reactions with over 10^9^ primers per bead (Fig. 1c). RT of *Gapdh* was not affected for DART-seq beads prepared with up to 10^10^ primers per bead.

Next, we evaluated the abundance of amplicons in sequencing libraries of reovirus infected cells generated by Drop-seq, and a DART-seq assay targeting all ten viral genome segments. We designed seven qPCR assays with amplicons distributed across two viral genome segments (S3 and L3). To account for assay-to-assay and sample-to-sample variability, we normalized the number of molecules detected in DART-seq and Drop-seq libraries to the number of *Gapdh* transcripts. We observed significant enrichment upstream (5’-end), but not downstream (3’-end) of the custom primer sites (Fig. 1d). Consistent with sequencing library preparation via tagmentation, we found that the degree of enrichment decreases with distance from the primer site.

We applied DART-seq to investigate the heterogeneity of cellular phenotypes and viral genotypes during T3D reovirus infection. Recent studies have explored RNA virus infection biology in single cells^4–6^, but were limited by cell throughput or restricted to the analysis of polyadenylated viral mRNAs. We infected L cells at a high multiplicity of infection (MOI 10), ensuring nearly all cells were infected (Fig. 2a). We performed Drop-seq and DART-seq experiments on infected and non-infected cells and implemented two DART-seq designs. The first targeted each viral genome segment with a single amplicon. The second comprised seven amplicons distributed across the S2 segment (Fig. 2b).

**Figure 2.**
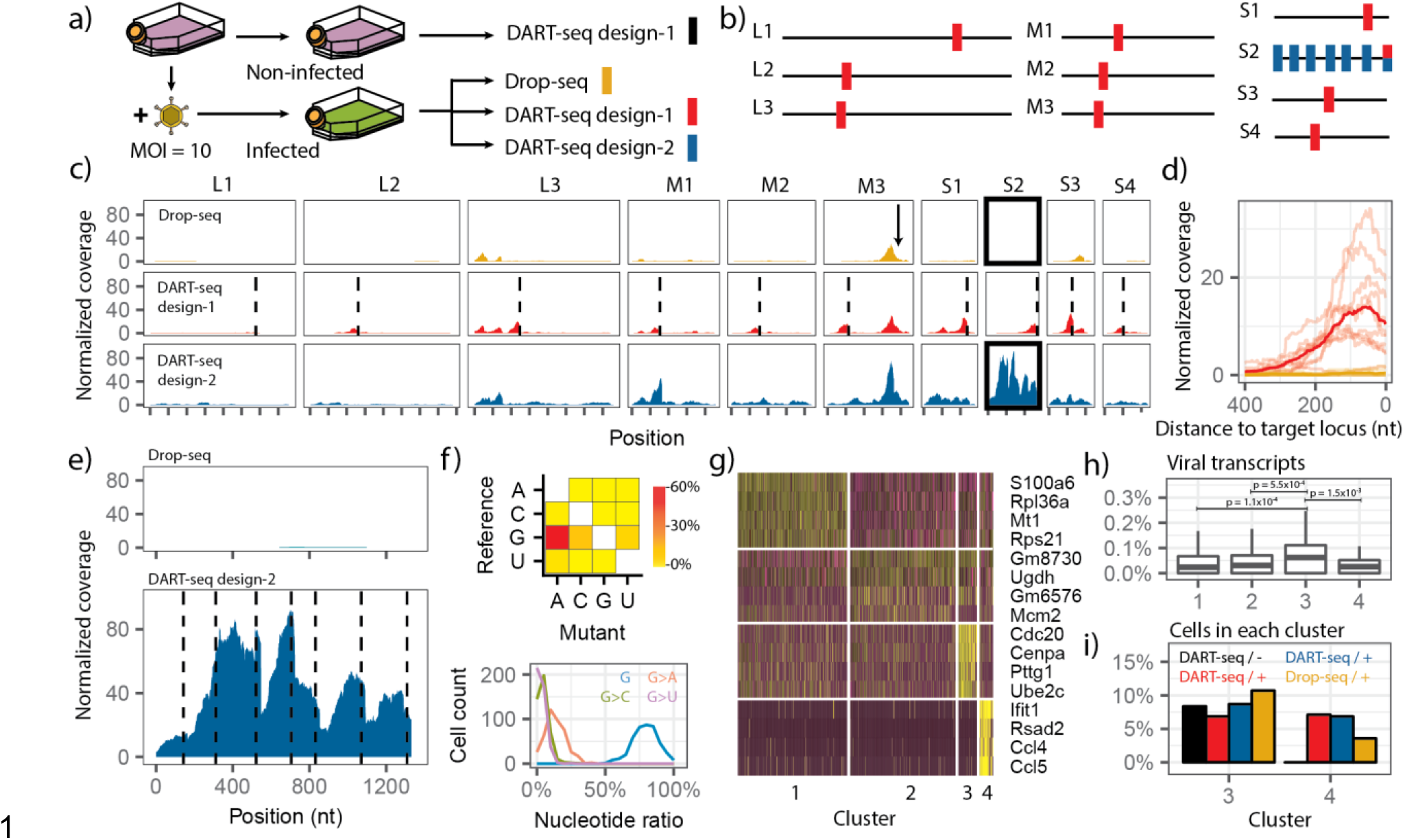
DART-seq reveals heterogeneity in viral genotypes and host response to infection. (a) Experimental design. (b) Schematic of DART-seq designs (design-1, red bars, design-2 blue bars). (c) Comparison of the sequence coverage (normalized to host UMI detected × 106) of the 10 reovirus gene segments (columns) for three different library preparations (rows). Arrow indicates an A5 pentanucleotide sequence part of segment M3. Dotted lines indicate DART-seq target positions. (d) Per-base coverage upstream (5’-end) of 10 custom primers of DART-seq design-1 (light red, average shown in dark red), and mean coverage achieved with Drop-seq (yellow). (e) Per-base coverage of the S2 gene segment achieved with DART-seq design-2 (bottom, dashed lines indicate custom primer positions) and Drop-seq (top). (f) Frequency and pattern of base mutations (top); histogram of nucleotide ratios for positions with reference nucleotide G detected in single cells (bottom). (g) Clustering analysis for variable gene expression of reovirus infected L cells (DART-seq design-1, yellow/purple is higher/lower expression). Similar clustering was observed in all three experiments with infected cells. (h) Relative abundance of viral transcripts in L-cell clusters (p-values determined by two-tailed Wilcox rank sum test). Lower and upper hinges correspond to 25th and 75th percentiles, respectively. Lower/upper whisker corresponds to smallest/largest value within 150% of the interquartile range from the nearest hinge (cluster 1, n = 411; cluster 2, n = 397; cluster 3, n = 50; cluster 4, n = 69). (i) Fraction of cells in meta-clusters for four experiments depicted in panel a with assay type and infection status (+ or -) indicated.

We analyzed the sequence coverage upstream of the DART-seq target sites. For both DART-seq designs, all targeted sites were enriched compared to Drop-seq (Fig. 2c, d). For design-1, we observed a mean enrichment of 34.7x 200 nt upstream of the custom primer sites. Viral transcripts were detected in Drop-seq upstream of A-rich sequences in the viral genome, consistent with spurious RT priming by oligo(dT) primers (Fig. 2c). Viral sequences were not detected in DART-seq or Drop-seq assays of non-infected cells. Experiments on an independent sample revealed similar sequencing coverage tracks across the reovirus genome (Fig. S3).

We tested the utility of DART-seq to measure the heterogeneity of viral genotypes in single cells with DART-seq design-2, which was tailored to retrieve the complete S2 segment. DART-seq design-2 increased the mean coverage across S2 430-fold compared to Drop-seq (cells with at least 1500 UMIs, Fig. 2e), enabling point mutation analysis. We identified 176 single-nucleotide variants (see Methods). Mutations from guanine-to-adenine (G-to-A) were most common (58%; Fig. 2f, top), though G-to-A mutational loads (mean 13%) varied across cells (Fig. 2f, bottom). We did not observe the G-to-A hypermutation in a highly expressed host transcript (*Actb*). The high G-to-A transition rate in viral transcripts could be secondary to a defect in viral transcription fidelity. The T3D strain used in this study has strain-specific allelic variation in polymerase co-factor μ2, which may affect the capacity of μ2 to associate with microtubules and the encapsidation of viral mRNAs^7,8^.

We identified four distinct cell subpopulations, after dimensional reduction and unsupervised clustering (design-1, Fig. 2g; Methods). Two major clusters comprised cells with elevated gene expression related to transcription and replication (*Rpl36a*, cluster 1) and metabolic pathways (*Ugdh*, cluster 2). Two additional clusters revealed upregulation of genes related to mitotic function (*Cdc20*, cluster 3) and innate immunity (*Ifit1*, cluster 4). The abundance of viral transcripts relative to host transcripts was significantly elevated for cells in cluster 3 (Fig. 2h; *p* = 1.0 × 10^−4^). We combined all datasets and quantified the cell type composition for each experiment. We did not observe cells related to cluster 4 (immune response) for the non-infected control (Fig. 2i).

We next explored the biological corollary of infection, the cellular immune response. The adaptive immune response relies on a diverse repertoire of membrane-bound and free antibodies. Antibody repertoires have previously been examined at depth^9,10^ but DART-seq widens the scope of such studies by providing concurrent transcriptome information. Antibodies are comprised of heavy and light chains, linked by disulfide bonds. Each chain contains variable and constant domains. The variable region is comprised of variable (V), diversity (D) and joining (J) segments in the heavy chain, and V and J segments in the light chain. We designed DART-seq primers to target VDJ and VJ gene segments in heavy and light chain transcripts^11^ (Fig. 3a).

**Fig. 3.**
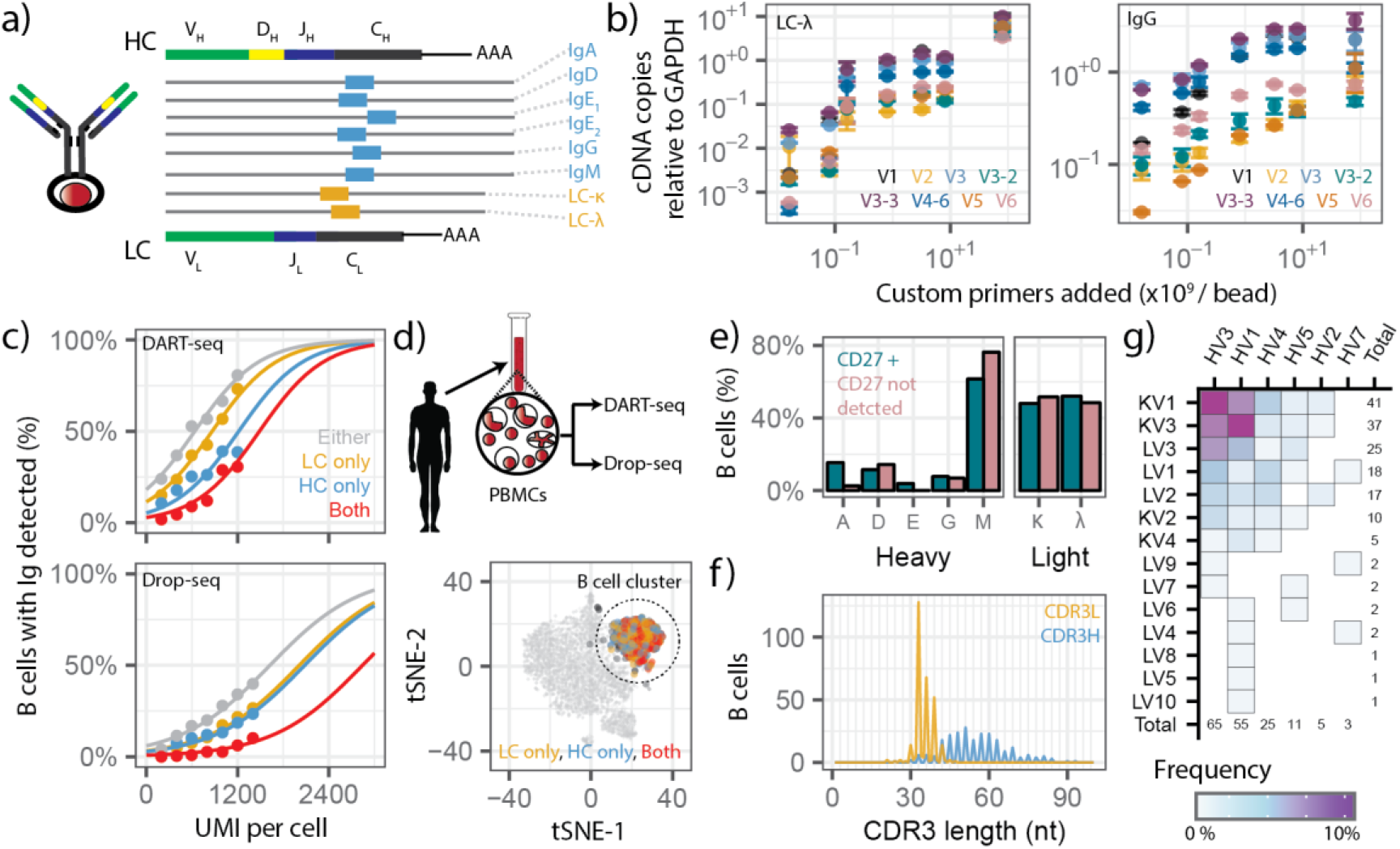
DART-seq measures paired heavy and light chain B cell transcripts at single cell resolution. (a) DART-seq custom primer design targeting the constant region of human heavy and light isotypes. (b) cDNA copies of Ig transcripts relative to *GAPDH* as a function of the number of custom primers included in the ligation reaction (left panel, LCλ+V primers; right panel: IgG+V primers, 62500 cells, 12000 beads, bulk assay). Points are mean of two replicate measurements, bars indicate the minimum and the maximum. (c) Percentage of B cells for which heavy and/or light chain transcripts were detected as a function of the UMI count per cell. Cells were binned by the number of UMI detected (bin width 200 UMI, 0-2400 UMI per cell, bins with fewer than 20 cells omitted, 26 - 2396 cells per bin). Distributions were fit with a sigmoid curve (Methods). (d) Drop-seq and DART-seq assays of human PBMCs. Experiments were performed on two distinct PBMC samples (n=2). Representative t-distributed Stochastic Neighbor Embedding (tSNE) for one DART-seq assay shown here (4997 single cells). Cells are colored based on heavy and/or light chain transcript detection. (e) Bar graph of isotype distribution for CD27+ B cells and B cells for which CD27 was not detected. (f) CDR3L and CDR3H length distribution. 818 B cells were used for the analysis. (g) Paired heavy (IGHV) and light (IGKV and IGLV) variable chain usage in B cells, (164 single cells).

We examined the efficiency of heavy and light chain transcript RT by qPCR (CD19+ B cells). We observed an enrichment of transcripts for all isotypes tested, as the number of custom primers on DART-seq beads was increased (Fig. 3b). Next, we compared the performance of DART-seq and Drop-seq to describe antibody repertoires (Fig. 3c). Approximately 120,000 B cells were loaded in each reaction, yielding 4909 and 4965 transcriptomes for DART-seq and Drop-seq, respectively. The number of UMIs and genes detected per cell was similar for DART-seq and Drop-seq (Fig. S4). We mapped transcript sequences to an immunoglobulin (Ig) sequence database (Methods). For both DART-seq and Drop-seq, the percentage of cells for which Ig transcripts were detected scaled with the number of UMIs detected in the cells (Fig. 3c). The Ig transcript recovery rate was significantly greater for DART-seq. For cells with 1000-1200 UMIs, we identified both heavy and light chain transcripts in 29% of cells using DART-seq, but in only 3% of cells using Drop-seq.

Next we applied DART-seq to determine the B cell antibody repertoire within human PBMCs (120,000 PBMCs, 4997 single-cell transcriptomes). To identify B cells, we used dimensional reduction and unsupervised clustering (Methods). We detected Ig transcripts in 564 of the 818 cells in the B cell cluster, and Ig expression mapped accurately onto the B cell population (Fig. 3d). DART-seq again outperformed Drop-seq in the recovery of antibody transcripts (Fig. S5). To test the reproducibility of DART-seq, we assayed an additional PBMC sample and observed similar Ig recovery rates (Fig. S5). We performed isotype distribution analysis on CD27+ B cells (Fig. 3e). As expected, CD27+ B cells were a mixed population of heavy chain isotypes, with IgM most frequently observed, followed by IgD and IgA^12^ (Fig. 3e). Kappa and lambda light chain isotypes were equally represented, as expected^13–15^ (Fig. 3e). B cells for which we did not detect CD27 were predominantly of the IgM isotype^16^ (Fig. 3e). B cells derive their repertoire diversity from the variable regions of their heavy (IGHV) and light chains^17^ (IGKV, IGLV). DART-seq captured a more diverse population of variable isoforms than Drop-seq (Fig. S6).

DART-seq can pair variable heavy and light chain transcripts in single cells. Out of 564 Ig transcript positive cells, we mapped the complete CDR3L in 339 cells and the complete CDR3H in 236 cells. The complete CDR3L+ CDR3H region was detected in 120 B cells. The number of VH and VL transcripts in single cells was correlated, as expected (Fig S7). The CDR3L and CDR3H length distributions had maxima around 30 and 50 nucleotides, respectively, as described previously^11,18^ (Fig. 3f). In line with previous reports, promiscuous light chain pairing was observed in 73.5% of the repertoires in CD27-B cells^18^. Finally, we measured clone specific pairing for the heavy (IGHV) and light chain variable regions (IGKV, IGLV) in 164 single B cells (Fig. 3g). The highest pairing frequency was observed between the most highly expressed heavy and light chain transcripts, consistent with previous reports^10,19^.

In conclusion, we have presented DART-seq an easy-to-implement droplet microfluidics technology to perform simultaneous RNA amplicon sequencing and transcriptome profiling in single cells. DART-seq enables a range of new biological measurements. We applied DART-seq to study the single-cell heterogeneity of RNA virus infection, thereby expanding on recent work that has demonstrated the utility to capture viral transcripts, including non-polyadenylated ones, together with transcriptomes in single cells at throughput^4–6^. DART-seq combines the ability to study multiple non-polyadenylated viral RNAs with the simplicity of droplet microfluidics. We furthermore applied DART-seq to determine the paired heavy and light chain repertoire in human B cells. A number of methods to reconstruct paired heavy-light chains from B cells have been described^10, 18–20^. DART-seq provides a tunable chemistry to simultaneously capture both heavy/light chains and the rest of the transcriptome, which is a key piece of information to elucidate the nature of the antibody-producing B cell.

## ACKNOWLEDGMENTS

We thank Peter Schweitzer and colleagues at the Cornell Biotechnology Resource Center (BRC) for help with sequencing assays. This work was supported by US National Institute of Health (NIH) grant 1DP2AI138242 to IDV and National Science Foundation Graduate Research Fellowship Program (NSF-GRFP) grant DGE-1144153 to PB.

## AUTHOR CONTRIBUTIONS

PB, MS, CGD, JSLP and IDV designed the study. PB, MS, SHK, MH, PML and MMH carried out the experiments. PB, MS, MFZW and IDV analyzed the data. PB, MS and IDV wrote the manuscript. All authors provided comments.

## ONLINE METHODS

### Step-by-step protocol

A detailed step-by-step protocol, including all reagents and primers used, is included as a supplemental file.

### Primer bead synthesis

Single-stranded DNA (ssDNA) primer sequences were designed to complement regions of interest. The probes were annealed to the complementary splint sequences that also carry a 10-12 bp overhang of A-repeats (Supplementary table). All oligos were resuspended in Tris-EDTA (TE) buffer at a concentration of 500 μM. Double-stranded toehold adapters^21^ were created by heating equal volumes (20 μL) of the custom primer and splint oligos in the presence of 50 mM NaCl. The reaction mixture was heated to 95°C and cooled to 14°C at a slow rate (−0.1°C/s). The annealed mixture of dsDNA probes was diluted with TE buffer to obtain a final concentration of 100 μM. Equal amounts of custom primer probes were mixed and the final mixture diluted to obtain the desired probe concentration (8.03 × 10^8^ custom primers per bead for reovirus DART-seq design-1 and B-cell DART-seq, and 4.01 × 10^9^ custom primers for reovirus DART-seq design-2). 16 μL of this pooled probe mixture was combined with 40 μL of PEG-4000 (50% w/v), 40 μL of T4 DNA ligase buffer, 72 μL of water, and 2 μL of T4 DNA Ligase (30 U/μL, Thermo Fisher). Roughly 12,000 beads were combined with the above ligation mix and incubated for 1 hr at 37°C (15 second alternative mixing at 1800 rpm). After ligation, enzyme activity was inhibited (65°C for 3 minutes) and beads were quenched in ice water. To obtain the desired quantity of DART-seq primer beads, 6-10 bead ligation reactions were performed in parallel. All reactions were pooled, and beads were washed once with 250 μL Tris-EDTA Sodium dodecyl sulfate (TE-SDS) buffer, and twice with Tris-EDTA-Tween 20 (TE-TW) buffer. DART-seq primer beads were stored in TE-TW at 4°C.

### Cell preparation

Murine L929 cells (L cells) in suspension culture were infected with recombinant Type 3 Dearing reovirus^22,23^ at MOI 10. After 15 hours of infection, the cells were centrifuged at 600 × g for 10 minutes and resuspended in PBS containing 0.01% BSA. Two additional washes were followed by centrifugation at 600 × g for 8 min, and then resuspended in the same buffer to a final concentration of 300,000 cells/mL (120,000 cells/mL in replicate experiment). Human CD19+ B cells or PBMCs were obtained from Zen-Bio (B cells: SER-CD19-F, PBMCs: SER-PBMC-F). Cells were washed three times with PBS containing 0.01% BSA, each wash followed by centrifugation at 1500 rpm for 5 min, and then resuspended in the same buffer. The cell suspension was filtered through a 40 μm filter and resuspended to a final concentration of 120,000 cells/mL.

### Single cell library preparation

Single cell library preparation was carried out as described2. Briefly, single cells were encapsulated with beads in a droplet using a microfluidics device (FlowJEM, Toronto, Ontario). After cell lysis, cDNA synthesis was carried out (Maxima Reverse Transcriptase, Thermo Fisher), followed by PCR (2X Kapa Hotstart Ready mix, VWR, 15 cycles). cDNA libraries were tagmented and PCR amplified (Nextera tagmentation kit, Illumina). Finally, libraries were pooled and sequenced (Illumina Nextseq 500, 20 × 130 bp). 2.6 × 10^7^ to 3.7 × 10^7^ sequencing reads were generated for the experiments described in Figure 2. 4.2 × 10^7^ to 6.8 × 10^8^ sequencing reads were generated for the experiments described in Figure 3.

### qPCR measurement of reverse transcription yield

80,000 L cells or 62,500 B cells were lysed in one mL of lysis buffer, and placed on ice for 15 minutes with brief vortexing every 3 minutes. After lysis and centrifugation (14,000 RPM for 15 minutes at 4°C), the supernatant was transferred to a tube containing 12,000 DART-seq beads. The bead and supernatant mixture was rotated at room temperature for 15 minutes and then rinsed twice with 1 mL 6x SSC. Reverse transcription, endonuclease treatment, and cDNA amplification steps performed as described above, with the exception that all reagent volumes were decreased by 80%. Following cDNA amplification and cleanup (following manufacturer’s instructions, Beckman Coulter Ampure beads), the total yield of cDNA was measured (Qubit 3.0 Fluorometer, HS DNA).

### qPCR measurements of amplicon enrichment in sequencing libraries

0.1 ng DNA from sequencing libraries was used per qPCR reaction. Each reaction was comprised of 1 μL cDNA (0.1 ng/μL), 10 μL of iTaq™ Universal SYBR® Green Supermix (Bio-Rad), 0.5 μL of forward primer (10 μM), 0.5 μL of reverse primer (10 μM) and 13 μL of DNAse, RNAse free water. Reactions were performed in a sealed 96-well plate using the following program in the Bio-Rad C1000 Touch Thermal Cycler: (1) 95°C for 10 minutes, (2) 95°C for 30 seconds, (3) 65°C for 1 minute, (4) plate read in SYBR channel, (5) repeat steps (2)-(4) 49 times, (6) 12°C infinite hold. The resulting data file was viewed using Bio-Rad CFX manager and the Cq values were exported for further analysis. Each reaction was performed with two technical replicates.

### Fluorescence hybridization assay

Roughly 6,000 DART-seq beads were added to a mixture containing 18 uL of 5M NaCl, 2 μL of 1M Tris HCl pH 8.0, 1 μL of SDS, 78 μl of water, and 1 μL of 100 μM Cy5 fluorescently labeled oligo (see Supplementary Table). The beads were incubated for 45 minutes at 46°C in an Eppendorf ThermoMixer C (15”, at 1800 RPM). Following incubation, the beads were pooled and washed with 250 μL TE-SDS, followed by 250 μL TE-TW. The beads were suspended in water and imaged in the Zeiss Axio Observer Z1 in the Cy5 channel and bright field. A custom Python script was used to determine the fluorescence intensity of each bead.

### Fluorescence hybridization assay to determine ligation efficiencies

Roughly 3,000 DART-seq beads were added to a mixture containing 18 uL of 5M NaCl, 2 μL of 1M Tris HCl pH 8.0, 1 μL of SDS, 78 μl of water, and 1 μL of 100 μM Cy5 fluorescently labeled oligo (see Supplementary Table). The beads were incubated for 45 minutes at 46°C in an Eppendorf ThermoMixer C (15”, at 1800 RPM). Following incubation, the beads were pooled and washed with 250 μL TE-SDS, followed by 250 μL TE-TW. The beads were suspended in 200 μl of DNAse/RNAse free water and transferred to a Qubit assay tube (ThermoFisher Scientific, Q32856). Qubit 3.0 Fluorometer was set to “Fluorometer” mode under the “635 nm” emission setting. The tube was vortexed briefly and placed in the fluorometer for immediate readout. Two additional vortexing and measurement steps were performed.

### Single cell host transcriptome profiling

We used previously described bioinformatic tools to process raw sequencing reads^1^, and the Seurat package for downstream analysis^24^. Cells with low overall expression or a high proportion of mitochondrial transcripts were removed. For clustering, we used principal component analysis (PCA), followed by k-means clustering to identify distinct cell states. t-stochastic neighborhood embedding^25^ (tSNE) was used to visualize cell clustering. For meta-clustering, host expression matrices from all four experiments were merged using Seurat^24^. Cells with fewer than 2000 host transcripts were excluded from the analysis in Figure 2. Cells with fewer than 100 unique genes detected were excluded from the analysis in Figure 3.

### Viral genotype analysis

Sequencing reads that did not align to the host genome were collected and aligned to the T3D reovirus genome^26^ (GenBank Accession EF494435-EF494445). Aligned reads were tagged with their cell barcode and sorted. The per-base coverage across viral gene segments was computed (Samtools^27^ depth). Positions where the per-base coverage exceeded 50, and where a minor allele with frequency greater than 10% was observed, were labeled as single nucleotide variant (SNV) positions. The frequency of SNVs was calculated across all cells. For the combined host virus analysis, the host expression matrix and virus alignment information were merged. The per-base coverage of the viral genome was normalized by the number of host transcripts. Cells with fewer than 1500 host transcripts were excluded from the analysis.

### Immunoglobulin identification and analysis

Sequences derived from B cells were collected and aligned to a catalog of human germline V, D, J and C gene sequences using MiXCR version 2.1.5^28^. For each cell, the top scoring heavy and light chain variable regions were selected for subtyping and pairing analyses (Fig. 3e and Fig. 3g).

### Sigmoidal fitting heavy/light chain capture

The mapping for the fractions of B cells containing heavy chains or light chains was fit with the following sigmoidal function:

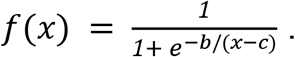

Where the parameter b was a free parameter for the fit of the light chain or heavy chain data, and then fixed for the light chain only, heavy chain only, and combined light chain and heavy chain data.

### Statistical analysis

Statistical tests were performed in R version 3.3.2^29^. Exact number of n values for each experiment are indicated in the figure legends. Error bars indicate the minimum and maximum of replicate measurements. Groups were compared using the two-tailed nonparametric Mann-Whitney U test. More information on the statistical parameters, sample size determination, and replication can be found in Life Sciences Reporting Summary.

## DATA AVAILABILITY

Raw sequencing data and corresponding gene expression matrices have been made available: NCBI Gene Expression Omnibus; Project ID GSE113675.

## CODE AVAILABILITY

Custom scripts are available at: https://github.com/pburnham50/DART-seq.

